# Random peptides rich in small and disorder-promoting amino acids are less likely to be harmful

**DOI:** 10.1101/2020.04.28.066316

**Authors:** Luke Kosinski, Nathan Aviles, Kevin Gomez, Joanna Masel

**Affiliations:** Department of Molecular and Cellular Biology, University of Arizona, Tucson, AZ 85721; Graduate Interdisciplinary Program in Statistics, University of Arizona, Tucson, AZ 85721; Graduate Interdisciplinary Program in Applied Math, University of Arizona, Tucson, AZ 85721; Department of Ecology and Evolutionary Biology, University of Arizona, Tucson, AZ, 85721

**Keywords:** Experimental evolution, evolvability, fitness estimation, preadaptation, de novo gene birth

## Abstract

Proteins are the workhorses of the cell, yet they carry great potential for harm via misfolding and aggregation. Despite the dangers, proteins are sometimes born *de novo* from non-coding DNA. Proteins are more likely to be born from non-coding regions that produce peptides that do little to no harm when translated than from regions that produce harmful peptides. To investigate which newborn proteins are most likely to “first, do no harm”, we estimate fitnesses from an experiment that competed *Escherichia coli* lineages that each expressed a unique random peptide. A variety of peptide metrics significantly predict lineage fitness, but this predictive power stems from simple amino acid frequencies rather than the ordering of amino acids. Amino acids that are smaller and that promote intrinsic structural disorder have more benign fitness effects. We validate that the amino acids that indicate benign effects in random peptides expressed in *E. coli* also do so in an independent dataset of random N-terminal tags in which it is possible to control for expression level. The same amino acids are also enriched in young animal proteins.

**Significance statement:** Proteins are sometimes born de novo. In an experiment to reproduce this process in *Escherichia coli*, we were able to predict 15% of the variation in random peptide fitness effects from their amino acid frequencies. In contrast, which order the amino acids are in seems to make no difference, adding no predictive power on top of simple amino acid frequencies. Amino acids that are smaller and promote intrinsic structural disorder have more benign fitness effects.

## Introduction

Proteins are the workhorses of the cell, but they are dangerous. For example, the polypeptide backbone is the key structural feature of amyloids, putting all proteins at risk of forming insoluble aggregates (Chiti and Dobson 2017), and most proteins are expressed at or just beyond their solubility limits (Vecchi, et al. 2020). Despite these dangers, new protein-coding genes are nevertheless born *de novo* from essentially random sequences (McLysaght and Guerzoni 2015; Van Oss and Carvunis 2019; Vakirlis, Carvunis, et al. 2020). To be beneficial enough for *de novo* birth, a random peptide must first do no serious harm, i.e. it must not be detrimental to the basic functioning of a cell. Here we quantify the degree to which, and the summary statistics via which, a random peptide’s propensity for harm can be predicted.

Neme et al. (2017) competed over 2 million *Escherichia coli* lineages, each containing a plasmid designed to express a unique random peptide, and tracked lineage frequencies over four days using deep DNA sequencing. This study has been criticized for providing too little support for the beneficial nature of the top candidates (Weisman and Eddy 2017; Knopp and Andersson 2018). But these criticisms do not detract from using the dataset to identify statistical predictors of serious harm versus relatively benign effect. Neme et al. (2017) used a strong promoter, so evaluation is of tolerance to high expression. Some fitness differences might be due to variation in expression e.g. due to auto-downregulation at the RNA level (Knopp and Andersson 2018) - we will return to this point in the last portion of the Results. Here we pursue analyses based on the hypothesis that the properties of the peptides contribute to variation in fitness among lineages.

Conveniently, computational predictors from peptide sequences alone are available for some properties, such as intrinsic structural disorder (ISD) and aggregation propensity. Because insoluble proteins have been implicated in toxicity and disease (Chiti and Dobson 2017) and peptides with high ISD are less prone to forming insoluble aggregates (Linding, et al. 2004; Angyan, et al. 2012), we hypothesize that highly disordered peptides are least likely to be strongly deleterious. Random sequences with high predicted disorder are well-tolerated *in vivo* (Tretyachenko, et al. 2017). Existing mouse (Wilson, et al. 2017) and Drosophila (Heames, et al. 2020) proteins, which are the product of evolution, are predicted from their amino acid sequences to be more disordered than what would be translated from intergenic controls.

Younger protein-coding sequences should be particularly constrained to first do no harm, as they have had little time to evolve more sophisticated harm-avoidance strategies (Foy, et al. 2019). In support of the idea that high ISD is an accessible way to avoid harm, young animal and fungal domains (James, et al. 2021) and genes (Wilson, et al. 2017; Foy, et al. 2019; James, et al. 2021), and novel overprinted viral genes (Willis and Masel 2018) have higher predicted disorder than their older counterparts. Some studies have found that putative de novo protein candidates in *Saccharomyces* yeasts have lower rather than higher ISD (Carvunis, et al. 2012; Basile, et al. 2017; Vakirlis, et al. 2018), but this could be an artifact of proportionately greater inclusion of non-genes, i.e. those for which there is no selection to retain a full-length translated peptide (Graur, et al. 2013), within the younger age classes. When Wilson et al. (2017) reanalyzed Carvunis et al.’s (2012) “proto-genes” of different ages, using more rigorous criteria to exclude non-genes from the data, the direction of the ISD trend was reversed. The same reversal of trend following a quality filter was also found by Vakirlis et al. (2018).

Protein ISD is determined both by the overall frequencies of the amino acids, and by the order in which those amino acids are arranged. Prior research on young genes has suggested that high predicted ISD in that context is driven primarily by amino acid frequencies, with amino acid order playing a more minor role (Wilson, et al. 2017). Here we ask the same question with respect to the role that both amino acid frequencies and their ordering have on peptide fitness, including through the promotion of ISD. Fortunately, the dataset of Neme et al. (2017) is large enough to look at the frequencies of each amino acid as predictors, rather than assume that existing prediction programs such as IUPred (Dosztányi, et al. 2005; Meszaros, et al. 2018) or Tango (Fernandez-Escamilla, et al. 2004; Linding, et al. 2004; Rousseau, et al. 2006) integrate all information about both amino acid frequencies and ordering in the best possible way. We can then test whether the way such programs integrate information about amino acid order gives them more ability to predict peptide fitness above and beyond the influence of amino acid frequencies. In doing so, we can estimate the relative roles of amino acid frequencies versus amino acid ordering in predicting fitness, as well as determine which amino acids have which effects.

Here we investigate the degree to which amino acid frequencies and amino acid ordering can predict the fitness effects of random peptides, and if so, which properties are most predictive. We also investigate whether the properties that help random peptides avoid harm in *E. coli* are also enriched in young eukaryotic proteins. With our work, we hope to further our understanding of how peptides avoid harm.

## Results

### Estimating the fitness effects of random peptides

Neme et al. (2017) tracked lineage frequencies over four days, and categorized a peptide as increasing or decreasing in frequency by comparing the DNA sequencing counts of day 4 to day 1 using DESeq2 (Love, et al. 2014). They report time information only in the form of days of the experiment, not the potentially non-constant number of generations. Even with this limitation, their approach fails to use the full richness of data on all 4 days, and applies a significance threshold that discards quantitative information.

We therefore reanalyze the same data, instead using a custom maximum likelihood framework (see Materials & Methods) to quantitatively estimate “fitness” and its associated confidence interval / weight. “Fitness” here refers to allele frequency changes over an entire cycle of population growth and dilution, rather than per generation. Our method classifies peptides quantitatively rather than qualitatively. It accounts for the fact that mean population fitness increases over the four days (see Materials and Methods). Our use of all available data within an appropriate maximum likelihood framework should make our method more sensitive and specific for identifying benign vs harmful peptides (see Supplementary Text).

Note that some peptides have such similar sequences that they should be considered pseudoreplicates (see Materials & Methods). We therefore grouped sequences into clusters based on sequence similarity (see Materials and Methods). There were 646 total clusters, of which there was statistical support for increases in frequency for the highest-weighted peptide in 138 clusters, and for decreases in 488 clusters. Some of our statistics use cluster as a random effect within a linear mixed model. To generate interpretable R^2^ values, we instead use a fixed effects model where we collapse each cluster into a single pseudo-datapoint with value given by the weighted mean and weight given by the sum of weights.

Due to the low number of sequences in some clusters, the residuals of our mixed model are subject to shrinkage (Savic and Karlsson 2009). We therefore present only results that remain significant in a fixed effect model in which we use only the highest weight peptide per cluster. These models are useful for ensuring our results are robust, but we do not use them as our primary model because they discard information, and so lose power on external sources of information (i.e. the correlations in Figs. 2, 3, and 6 are weaker in the fixed effects model).

### Most predictive power stems from amino acid frequencies rather than amino acid order

We estimated peptide disorder using several metrics that contain information both about amino acid frequencies and about their order: IUPred as an estimate of intrinsic structural disorder (Dosztányi, et al. 2005; Meszaros, et al. 2018), CamSol as an estimate of water solubility (Sormanni, et al. 2015), and Tango as an estimate of general aggregation propensity (Fernandez-Escamilla, et al. 2004; Linding, et al. 2004; Rousseau, et al. 2006). Fewer than 6% of the random peptides have a predicted transmembrane helix (Dataset S1) from TMHMM (Krogh, et al. 2001), so our choice of these predictors is guided by our assumption that the random peptides are predominantly located in the cytosol. Having a predicted transmembrane helix did not in itself predict random peptide fitness effects (P = 0.2, likelihood ratio test relative to mixed model with only the intercept as a fixed effect). In contrast, each of our cytosol-solubility-inspired metrics significantly predicted random peptide fitness (Fig. 1A – 1C), with effects in the predicted direction (more disorder and more solubility are good, more aggregation propensity is bad). Adjusted R^2^ values for IUPred, CamSol, and Tango are 0.029, 0.029, 0.016, respectively. Another aggregation predictor, Waltz (Maurer-Stroh, et al. 2010), that specializes in β aggregates, was also in the predicted direction of aggregation being harmful, but did not meet statistical significance (P = 0.07).

**Fig. 1.**
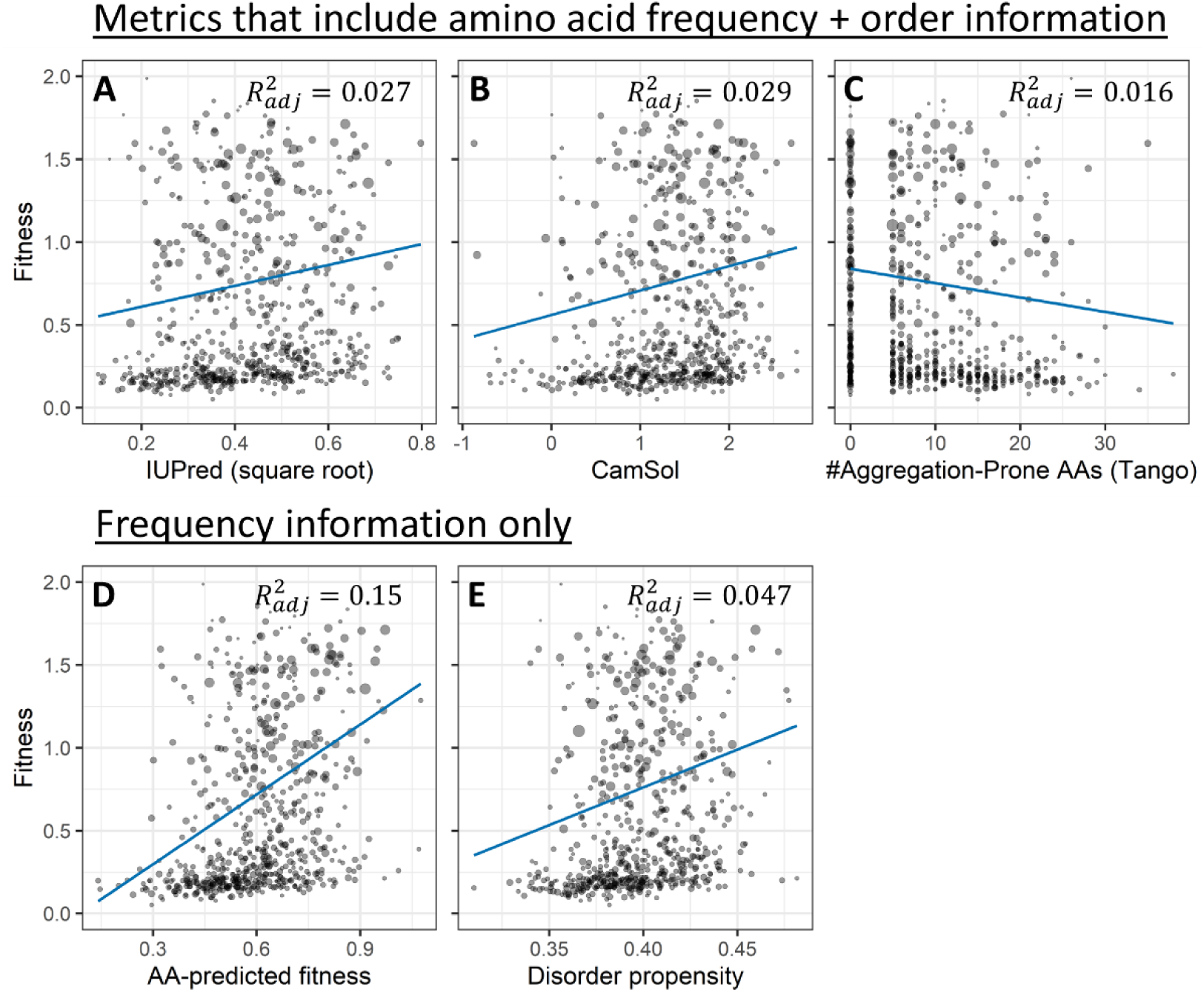
Many metrics predict peptide fitness effects, but most predictive power comes from amino acid frequencies. Three metrics that combine information on both amino acid frequencies and amino acid order ((A) IUPred, (B) CamSol, and (C) Tango), and two that contain only amino acid frequency information ((D) 19 custom weights on amino acid frequencies and (E) independently estimated disorder propensities used as weights on amino acid frequencies), each significantly predict peptide fitness on their own (P = 7 × 10^−4^, 0.003, 0.01, 5 × 10^−6^, and 9 × 10^−7^, respectively, likelihood ratio test in mixed model compared to intercept-only model). Each point (n = 646) shows a cluster of sequences with similar amino acid sequences (see Methods for more details), and the area displayed for each point is proportional to summed weights across that cluster. Blue lines are fixed-effect weighted linear regressions of cluster fitness on the x-axis predictor, where clusters are collapsed to a single pseudo-datapoint by their weighted average and weights are sums within each cluster. Metrics that include both frequency and order information fail to outperform frequency-only based metrics, as shown by regression slopes (blue lines) and adjusted R^2^ values (top right of each figure panel). Adjusted R^2^ is calculated as 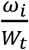, where *n* is the number of data points and *p* is the number of degrees of freedom in the predictor. Note that in part D the predictor (model-predicted fitness) is a composite of 19 degrees of freedom that have all been trained on the dataset, so care should be taken in comparing its blue regression line to that of the other panels, each of which has a predictor with only one degree of freedom – this problem does not apply to comparisons of adjusted R^2^ values. Seven clusters with fitness greater than 2 are not shown here for ease of visualization; a complete y-axis is shown in supplemental fig. 1. Log-transforming fitness would remove high fitness skew, but creates systematic heteroscedasticity, and so was not done (supplemental fig. 2). The lack of systematic heteroscedasticity can be seen here in the form of similar point size across fitness values.

Next we asked whether these sophisticated metrics offer additional predictive power beyond mere amino acid frequencies, in the light of prior work on young genes in which little additional predictive power was found (Wilson, et al. 2017). To do this, we fit a model of fitness predicted by amino acid frequencies, measured from counts of each amino acid in each peptide’s random region (Fig. 1D), and compared its performance to predictors that incorporate ordering information (Figs. 1A-C). The amino acid frequency-only model was a significant predictor of fitness (P = 2 × 10^−6^, likelihood ratio test compared to an intercept-only mixed model). It is also more biologically predictive than other metrics, with adjusted R^2^ = 0.15 (adjusted to account for the number of predictors used) being far greater than the values of 0.027, 0.029, and 0.016 found in Figs 1A-1C. Another, non-adjusted, way to look at biological effect size is the far steeper blue line in Fig. 1D than in Figs. 1A-1C. Statistically, when the frequencies of each of the twenty amino acids are used as predictors (Fig. 1D), then IUPred, CamSol, and Tango drop out of the model (P = 0.2 for each, likelihood ratio test in mixed model, see Supplemental Table S1), suggesting that their predictive power in Figs. 1A-1C came largely from being metrics of amino acid frequencies. These results are surprising: one might expect sophisticated metrics that incorporate both amino acid frequencies and order information to offer more predictive power and explain a greater range of fitness than simple amino acid frequencies, yet they fail to do so.

Our Fig. 1D model using the frequencies of the 20 amino acids involves 19 degrees of freedom, while the other metrics we examine involve only one. This makes it inappropriate to compare the slopes of the blue lines, although adjusted R^2^ values can still be compared, and the fact that the other metrics drop out of a combined model is also informative. We also investigated a one degree of freedom model of amino acid frequencies, in which relative weights were specified in advance by a disorder propensity metric that assigns each amino acid a score based on how frequently it is found in known disordered versus ordered proteins (Theillet, et al. 2013). Average disorder scores over each peptide’s random region significantly predicted random peptide fitness effects in a linear mixed model (Fig. 1E, P = 9 × 10^−7^, likelihood ratio test compared to an intercept-only model). The effect size on predicted fitness from the 10% to the 90% quantiles of disorder propensity is 0.50 to 0.72, and the adjusted R^2^ for the disorder propensity model is 0.047. For comparison to other predictors with a single degree of freedom, the model that got the largest effect size from incorporating both amino acid frequency and order information was IUPred with an effect size from 0.52 to 0.70, and CamSol had the highest adjusted R^2^ at 0.029. This superiority of the one degree of freedom disorder propensity model further suggests that predictive power resides with amino acid frequencies, not order information.

We next investigated a metric of ISD that is comprised of only order information. This can be calculated as the excess IUPred score of the real peptide in comparison to the average IUPred score of a set of hypothetical peptides in which the order of the amino acids has been randomly scrambled; this metric was previously found to be elevated in younger mouse genes (Wilson, et al. 2017). However, adding this ΔISD metric to our model with amino acid frequencies as predictors did not significantly improve the model (P = 0.2). This further supports our conclusion that amino acid ordering plays only a minor role compared to amino acid frequencies in the fitness effects of the random peptides examined here.

### Small and disorder-promoting amino acids predict benign fitness effects

Next we quantify the statistical effect of each of the 20 amino acids on fitness. Naively, we could take the associated slope coefficient in a multiple regression model, which represents the change in fitness when one amino acid is gained. But in a peptide of fixed length, one amino acid cannot be gained without another amino acid being lost. We therefore instead calculate the marginal fitness effect of each amino acid on fitness (see supplementary text and Table S2, displayed in fig. 2, y-axis), representing the effect of gaining that amino acid and losing a randomly selected alternative.

**Fig. 2.**
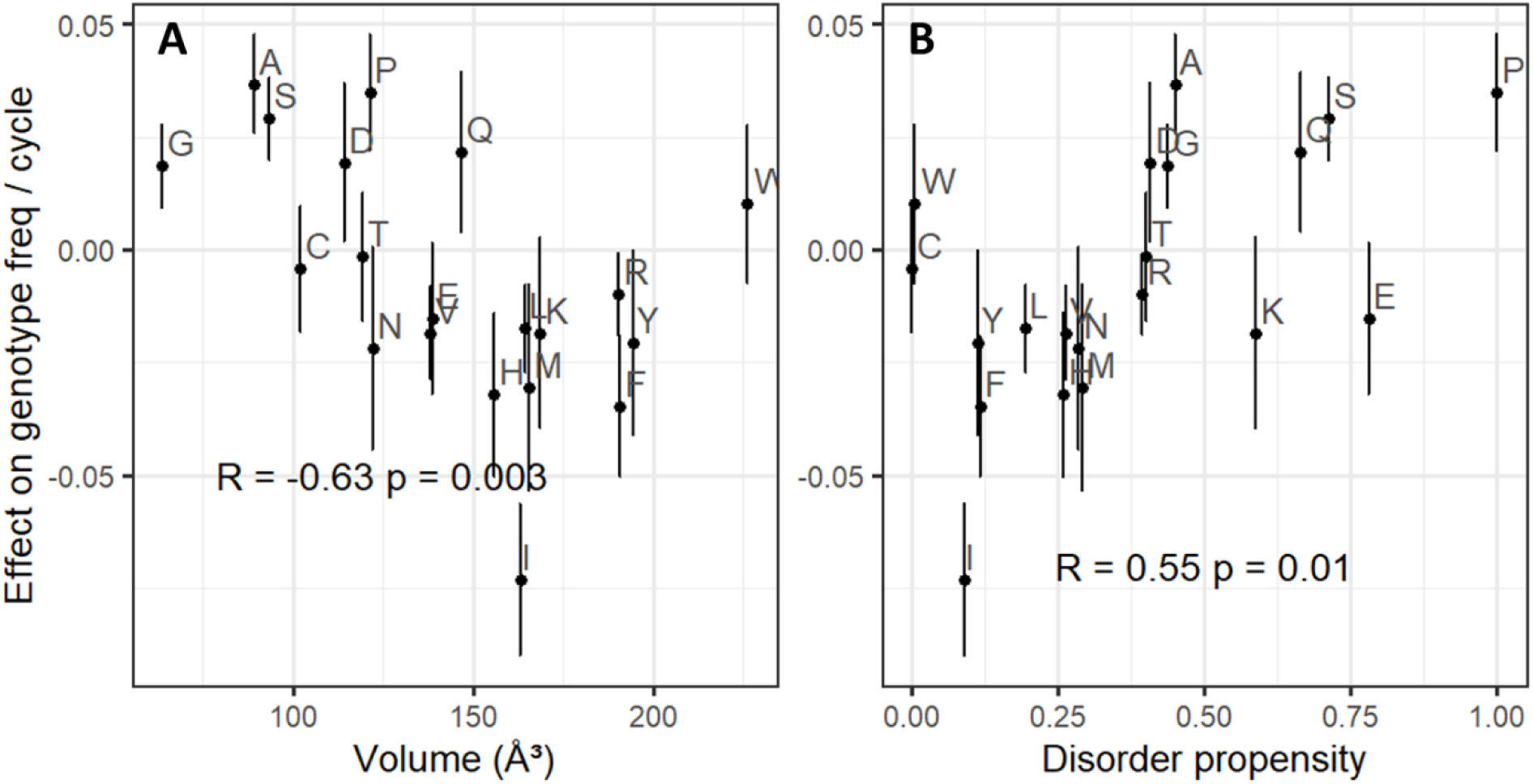
Amino acids that are small and are associated with disorder promote higher fitness. The y-axis shows each amino acid’s marginal effect on fitness, which is the change in fitness when one amino acid of the focal type replaces one randomly chosen amino acid of a different type in a random peptide (see Supporting Information). Error bars are +/-one standard error. P-values and correlation coefficients come from weighted Pearson’s correlations, where weights for marginal effects are calculated as 1 / s.e. (marginal fitness effect)^2^, and volume and disorder propensity are unweighted.

Amino acids with smaller volumes (Tsai, et al. 1999) and higher disorder propensities (Theillet, et al. 2013) tend to have higher marginal fitness effects (fig. 2A and 2B; P = 0.01 for both disorder propensity and volume, likelihood ratio test for dropping either term from a weighted regression of marginal effect on both volume and disorder propensity). Volume and disorder propensity together explain over half the weighted variation in marginal fitness effect (weighted adjusted R^2^ = 0.52). Other properties of amino acids, such as stickiness (Levy, et al. 2012), relative solvent accessibility (Tien, et al. 2013), amino acid cost in *E. coli* (Akashi and Gojobori 2002), frequency in the *E. coli* proteome (either raw or the excess/deficiency expected from number of codons; see Materials and Methods), and isoelectric point (Liu, et al. 2004) did not provide significant explanatory power on top of disorder propensity and volume (all P > 0.1, likelihood ratio test).

Tryptophan is an outlier for amino acid effects on fitness, with a slightly positive effect on fitness despite both its large volume and its underrepresentation in disordered regions (fig. 2). Removing tryptophan from a weighted regression model of volume and disorder propensity predicting marginal effect increases the weighted adjusted R^2^ from 0.52 to 0.68. Tryptophan, encoded only by UGG, is nearly 60% more common among peptides with at least 5 sequence reads than we expect from the 58% GC content of our dataset. Together with the confidence interval for its marginal fitness effect including 1, this provides further evidence that tryptophan is not harmful, making it a distinct outlier, for reasons that are not clear to us.

Isoleucine also stands out, as even more harmful than expected by its large size and order propensity. Isoleucine’s harmful effects may be exacerbated by its role in amyloid formation. For example, familial amyloid cardiomyopathy is most commonly caused by a valine to isoleucine mutation (Jacobson, et al. 1997; Dubrey, et al. 2015), suggesting that isoleucine has potential to form dangerous amyloids where other hydrophobic amino acids do not. Isoleucine, valine, and leucine are all hydrophobic amino acids with a branched carbon, but only raised isoleucine levels are associated with a higher risk of Alzheimer’s disease (Larsson and Markus 2017), further suggesting that isoleucine may be especially prone to amyloid formation.

### Young animal sequences are enriched for amino acids that increase fitness in random peptides

As discussed in the Introduction, young domains have higher predicted ISD than their older counterparts. One hypothesis to explain this observation is that in order to be successfully born *de novo*, a protein sequence is especially constrained to first do no harm (Wilson, et al. 2017). However, the “phylostratigraphy” approach of assigning ages to genes, on the basis of the species range in which they have homologs, is contentious. Detecting homologs is more difficult for fast-evolving sequences, which may be erroneously scored as young (Alba and Castresana 2007; Moyers and Zhang 2015, 2016). Disordered proteins tend to be fast evolving (Chen, et al. 2011), suggesting that highly disordered genes could be misclassified as young because of their fast evolutionary rate. If the amino acid enrichments of higher fitness random peptides match the amino acid enrichments of young genes, this would be evidence that the *de novo* gene birth process, rather than homology detection bias alone, causes trends in protein properties as a function of apparent gene age.

To test this, we took the slopes of amino acid frequencies with protein domain age from James et al. (2021), as quantified across over 400 eukaryotic species. As predicted, amino acids that are good for random peptides are enriched among the youngest animal Pfams (fig. 3A). This prediction was not, however, supported for trends among recent plant domains (fig. 3B) nor among ancient (fig. 3C) domains older than 2.1 billion years. Plant and ancient trends reflect a *de novo* gene birth process that enriches for the most abundant amino acids in their respective lineages, such as cysteine, rather than for amino acids that promote ISD (James, et al. 2021). It is interesting that we find that ISD still predicts harmlessness in *E. coli*, even though we do not find evidence it shaped *de novo* gene birth in its distant ancestors. We also note that ISD does shape recent *de novo* gene birth in viruses (Willis and Masel 2018).

**Fig. 3.**
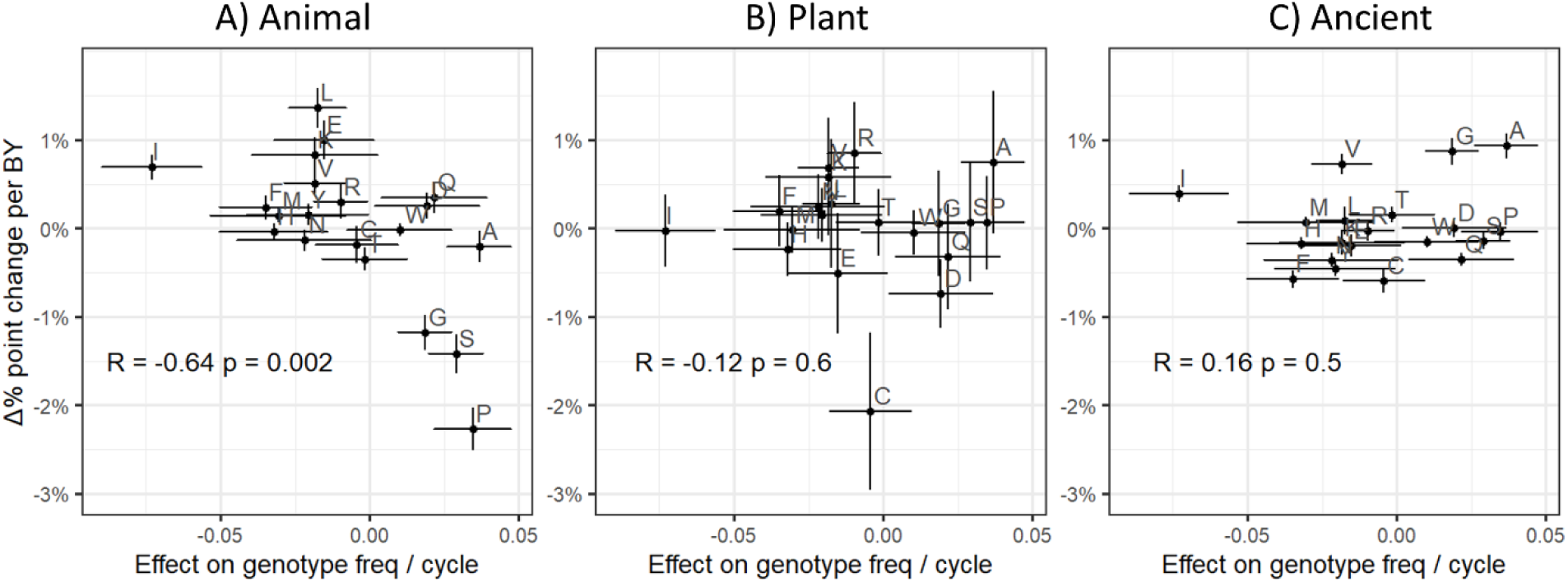
Purportedly young animal Pfams are enriched for amino acids that predict high fitness in random peptides. The y-axis represents how the frequency of each amino acid depends on the age of the sequence in billion years (BY), estimated as a linear regression slope for non-transmembrane Pfam domains (James, et al. 2021). Frequency is in number of percentage points, e.g. a difference in glutamic acid content of 5% vs. 6% is a difference of one percentage point. The x-axis shows each amino acid’s marginal effect on fitness, which is the change in fitness when one amino acid of the focal type replaces one randomly chosen amino acid of a different type in a random peptide (see Supporting Information). Error bars are +/- one standard error. Fitness effects predict A) animal, but not B) plant, or C) ancient (older than 2.1 billion years) Pfam phylostratigraphy slopes. Correlation coefficients and P-values come from weighted Pearson correlations. Note that the P-value for animal phylostratigraphy slopes vs marginal effects survives a conservative Bonferroni correction (P = 0.002 < 0.05/3 = 0.017).

### Fitness is better predicted by amino acid frequencies than by GC content

Long et al. (2018) proposed that selection acts directly on GC content, perhaps due to the three hydrogen bonds of G-C pairs. Amino acids encoded by Gs and Cs tend to promote higher ISD (Angyan, et al. 2012), making it difficult to distinguish between selection for high GC content and selection against harmful amino acids. To attempt to distinguish between the two, we compare amino acids that always have G or C to those that always have A or T, at both the first and second nucleotide positions in the codon. If selection were for GC nucleotides, we would expect GC to predict high marginal amino acid fitness effects at both positions. But if results are dramatically different at the two positions, this would show that it is selection on amino acid content that drives GC as a correlated trait. Results are statistically significant in the predicted direction at the second position (fig. 4A, P = 0.001, weighted Welch’s t-test), and in the predicted direction but not statistically significant at the first (fig. 4B, P = 0.2). The effect size of GC content on fitness could not be statistically distinguished between the first and second position (fig. 4C), with wide and hence inconclusive error bars.

**Fig. 4.**
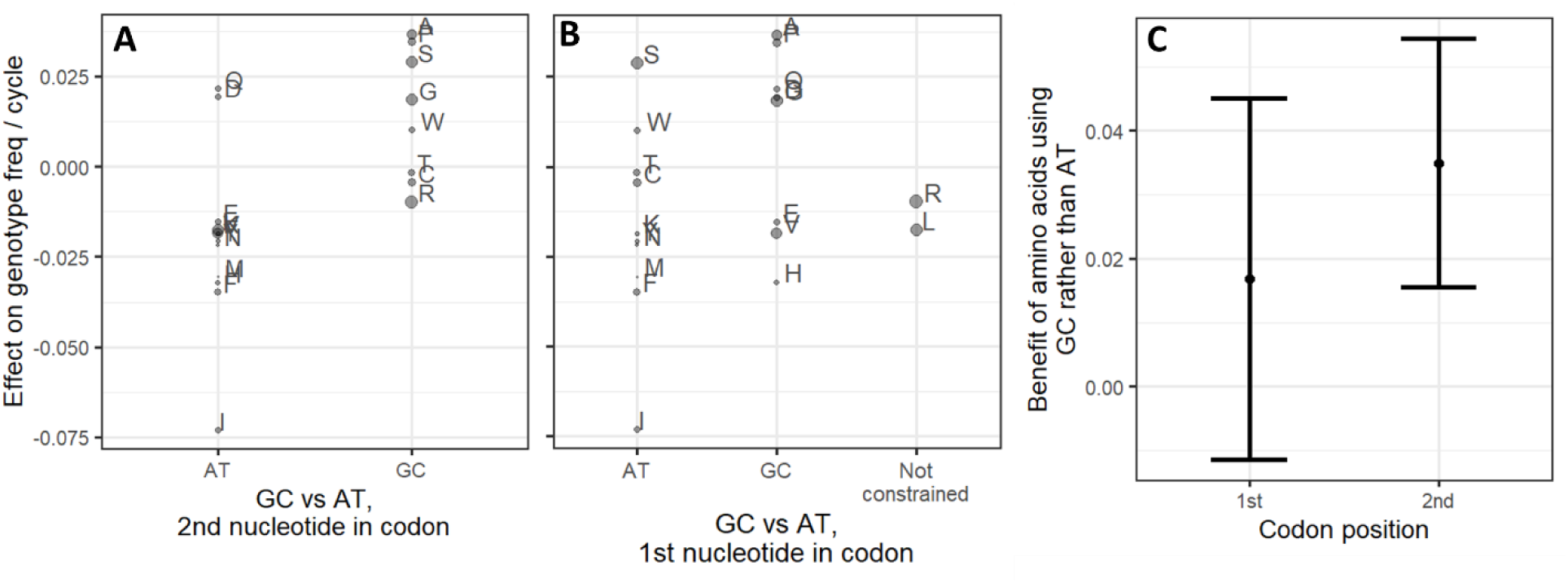
Amino acids that are constrained to use Gs and Cs tend to have higher marginal effects on fitness than those constrained to use As and Ts. The difference is significant for constraints at the second nucleotide position of a codon (A) (P = 0.001, weighted Welch’s t-test), but not at the first (B) (P = 0.2). Point area is proportional to weight, which is calculated as 1 / s.e.(marginal fitness effect)^2^, as described in Supporting Information. The y-axis is the same as the fig. 2 y-axis and fig. 3 x-axis. C) The mean advantage of amino acids constrained to use GC rather than constrained to use AT is not distinguishable in size between the first and second codon positions. Y-axis gives the difference in the two weighted means of marginal fitness effects from A) and B). Error bars represent 95% confidence intervals on the difference between the means (calculated as difference +/-t_crit_ × se), where t_crit_ ≈ 2.1 is the critical value of the t-statistic with the appropriate degrees of freedom. Weighted Welch’s t-test statistic and the corresponding standard error of the difference in means were calculated using the “wtd.t.test” function from the “weights” R package, version 1.0.1.

Linear models are compatible with partially independent contributions of both amino acid frequencies and GC content to harm avoidance. GC content, calculated from the random portion of each peptide’s sequence (for more details, see Materials and Methods), is a statistically significant predictor of fitness by itself (P = 6 × 10^−11^, likelihood ratio test for nested fixed-effect models relative to intercept-only model). However, the weighted adjusted R^2^ of 0.06 for GC content is much lower than the weighted adjusted R^2^ of 0.15 for full amino acid frequency information, i.e., it explains less of the variation than amino acid frequencies. Adding GC content to the amino acid frequencies-only model offers only a modest improvement (P = 0.003, weighted adjusted R^2^ values improves from 0.15 to 0.16), while adding amino acid frequencies to a GC content only model offers a notably larger improvement (P = 1 × 10^−11^, weighted adjusted R^2^ improves from 0.06 to 0.16). These weighted adjusted R^2^ values suggest that while there may be some direct selection on GC content, the effect of amino acid frequencies appears to be well beyond what can be explained by GC content.

To further verify that GC content is not the primary driver of our results, we crudely controlled for %GC by splitting our dataset into high (> 57.4%) and low (≤ 57.4%) GC random sequences and repeated the analysis of figure 1D for each subset. The 57.4% cutoff was the median GC among the pseudo-datapoints corresponding to clusters. High and low %GC data subsets produced nearly identical fits to each other (fig. 5) and to figure 1D. The adjusted R^2^ is 0.11 for the high and 0.09 for the low GC content subsets, with the drop to be expected given the restriction of range. If %GC were a major driver, we would expect an offset between the regression lines in fig. 5. Instead, these results are compatible with amino acid frequencies being the primary driver of results, with %GC being mostly just a correlated trait with little causal explanatory power.

**Fig. 5.**
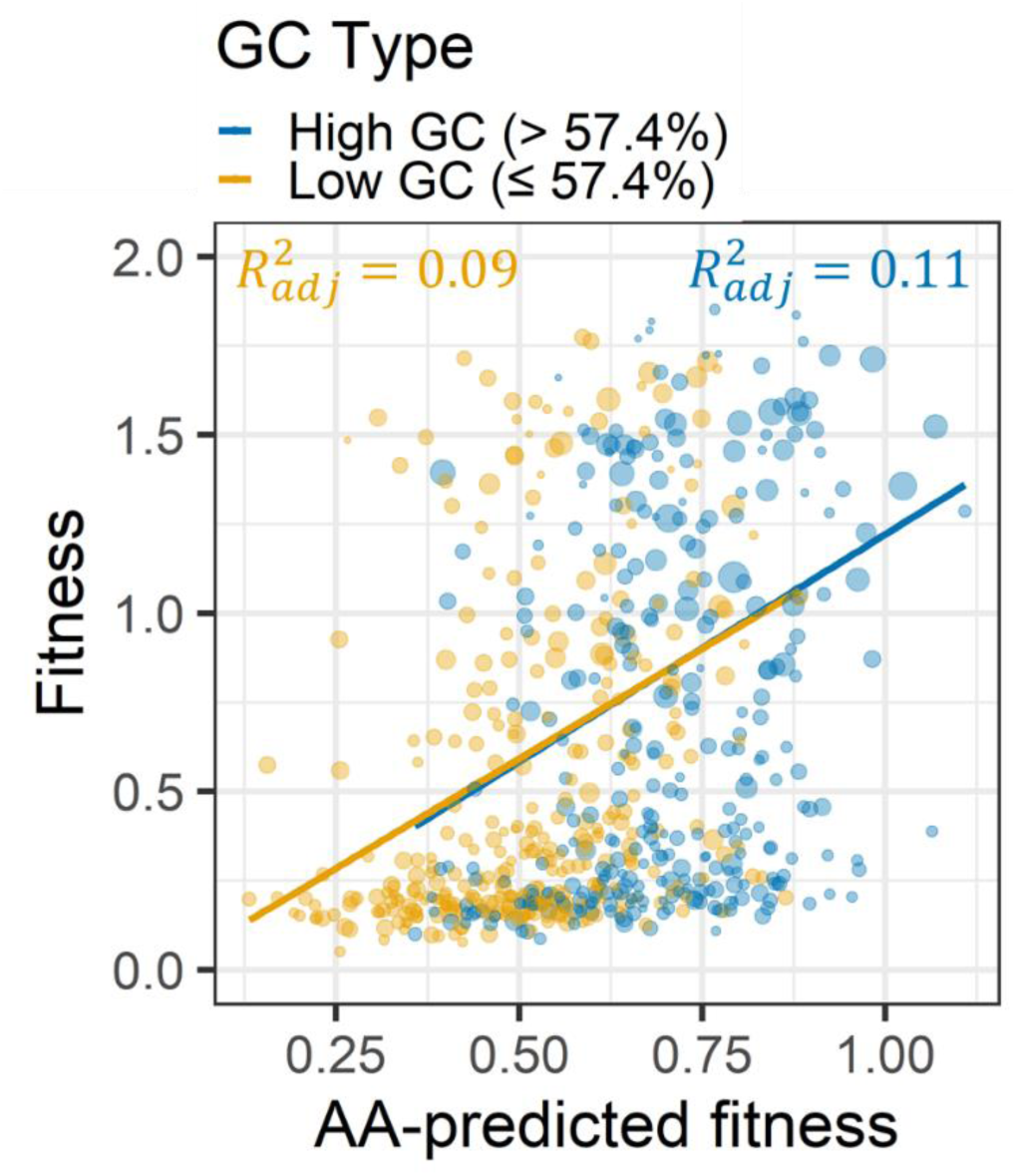
Amino acid frequencies predict fitness in the same way for peptides encoded by high vs. low GC content sequences. We split data into high (> 57.4%) and low (≤ 57.4%) GC content and separately fit a model to each in which amino acid frequencies predicted peptide fitness, as in figure 1D. Statistical significance is best assessed with GC content as a quantitative rather than a binary predictor (described in text). For the full methodological details of this figure, please refer to the Figure 1 legend.

### The same amino acids predict benign fitness effects in random N-terminal tags

The degree to which benign effects are due to low expression of a random peptide, vs. benign effects of the peptide once expressed, remains unclear. We therefore tested the ability of our amino-acid-frequencies-only model, trained on the data of Neme et al. (2017), to predict residual fitness effects in a dataset that controls for peptide expression level. Goodman et al. (2013) tagged the N-prime end of green fluorescent protein (GFP) with 137 different short random sequences (11 amino acids long), allowing random peptide expression level to be measured via fluorescence. Frumkin et al. (2017) measured the fitness effects of these random peptide-tagged GFPs in *E. coli* using FitSeq (Li, et al. 2018). For 89 of them, Frumkin et al. (2017) were able to calculate a “fitness residual” based on the deviation from the fitness expected from the level of GFP expression. Note that while this fitness residual controls for expression level, it still contains the cost of inefficient expression in addition to the fitness effect of the peptide itself. Frumkin et al. (2017) found that low fitness residuals were associated with hydrophobic and expensive-to-synthesize amino acids. These findings are consistent with our own estimates of direct peptide effects, as hydrophobic amino acids tend to be order-prone (Linding, et al. 2004; Angyan, et al. 2012), and amino acid volume is highly correlated with synthesis cost in *E. coli* (Pearson’s correlation coefficient = 0.85, P = 2 × 10^−6^, cost for amino acid synthesis in *E. coli* taken from (Akashi and Gojobori 2002)). Indeed, predicted fitness values for Frumkin et al.’s (2017) N-terminal tags were significantly correlated with their actual fitness residuals (fig. 6). The consistency between our results and the findings of Frumkin et al. (2017), who control for peptide expression level, provides an external validation of our results and suggests that our findings are unlikely to be due to differences in peptide expression levels alone.

**Fig. 6.**
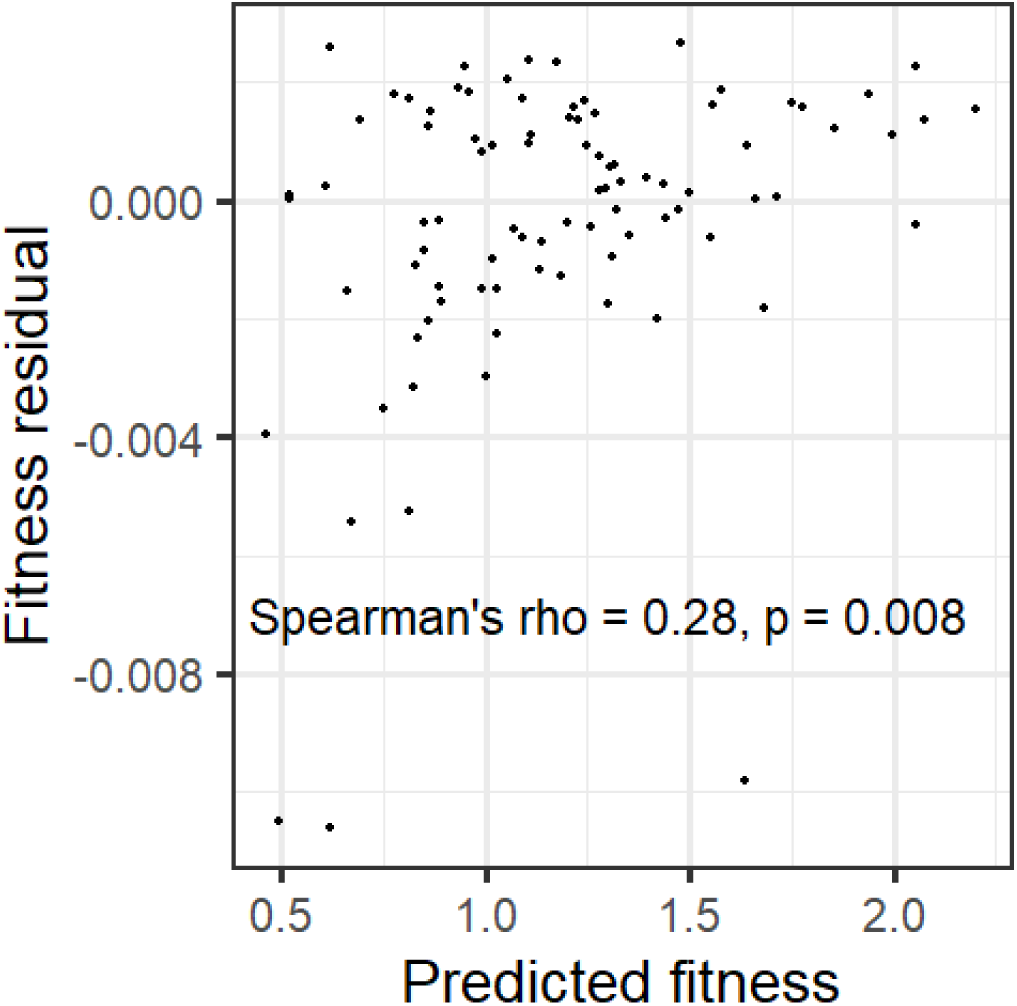
Fitness predictions trained on the random peptides of Neme et al. (2017) also work for short random tags attached to the N-terminus of GFP. Predicted fitness comes from our amino acid frequencies-only mixed model. “Fitness residuals” of N-terminal tags are from Frumkin et al. (2017), and represent the difference between the fitness of the construct and the expected fitness from expression level. *n* = 89.

## Discussion

We found that, while many metrics of peptide properties have some ability to predict the fitness effects of random peptides expressed in *E. coli*, most predictive power stems from amino acid frequencies. Simply knowing how many of which amino acids are present in these random peptides can account for 15% of the variance in fitness among lineages, and adding more predictors to account for amino acid order fails to add more predictive power. This indicates both the success of our statistical method for inferring fitness, and that mere amino acid frequencies without amino acid order can be informative of peptide fitness effects. Amino acids that are small and promote disorder predict high fitness in *E. coli*, and align with those that are enriched in young protein domains in animals.

Most studies of random peptides have focused on finding peptides that have specific binding or function (e.g. Kaiser, et al. 1987; Keefe and Szostak 2001; Frulloni, et al. 2009). Some were motivated as proof-of-concept that random peptides can exhibit properties of native proteins, such as folding (Davidson and Sauer 1994; Chiarabelli, et al. 2006; LaBean, et al. 2011) and being soluble (Prijambada, et al. 1996). Others focus on how to increase the percentage of native-like random peptides, e.g. by showing that more hydrophilic random peptide libraries have a higher percentage of stable and soluble peptides (Davidson, et al. 1995). Our work has a different intent, identifying properties that make a peptide less likely to be harmful. Neme et al.’s (2017) experiment was suitable for this purpose because it used a large library of peptides with diverse properties, competed lineages growing under permissive conditions, and measured relative growth rates (i.e. fitness). In contrast, a study design such as that of Knopp et al. (2019), who selected random peptides that rescue viability in the presence of antibiotics, is less suitable for our purposes because so few peptides, including harm-avoiding peptides, are viable. Neme et al.’s (Neme, et al. 2017) study was also convenient because all peptides were the same length – 65 amino acids with 50 amino acids of random sequence – allowing us to neglect length in our analysis (see (Castro and Tautz 2021) for the analysis of shorter sequences).

Having a higher proportion of random peptides do no harm is expected to increase the success rate of future screens for peptides with specific properties. Nucleotide sequences with high %GC content tend to encode peptides with more benign fitness effects, suggesting that higher %GC should be used in future random peptide libraries. However, very high GC content will yield low complexity sequences, which our predictor has not been trained on. The marginal fitness effects of each amino acid might be different in this very different context.

While the library used by Neme et al. (2017) was designed to have equal frequencies of each nucleotide in the random region, and thus 50% GC content, the over two million random peptides that had at least one sequencing read had a GC content of ∼59% in their random portion. The mean GC content of the peptide clusters we analyzed (see Materials and Methods) was similar, at ∼58%, with higher fitness peptides within this group having still higher %GC, as discussed in the Results. The enrichment from 50% GC to ∼59% GC might be because many lower GC content sequences were so harmful that lineages that carried them went extinct prior to detection via sequencing. Note that sequencing methods vary in their GC bias (Benjamini and Speed 2012; Choudhari and Grigoriev 2017), making these absolute values less reliable, and making it important to use internal controls. Our use of four time points acts to some degree as such an internal control, except to the degree that bias changes with %GC over the course of the experiment.

Long et al. (2018) proposed that there is direct selection for high GC content, as evidenced in part by a preference for amino acids with G or C at the second position of codons, in excess of that predicted from mutation accumulation experiments. Our findings cannot completely exclude this hypothesis, but show stronger selection on amino acid frequencies, selection that is capable of driving increased GC content in coding regions as a correlated trait. In intergenic regions, elevated %GC is likely driven mostly by GC-biased gene conversion. However, elevated GC content could also be due, at least in part, to selection on peptides from non-coding regions translated by error (Rajon and Masel 2011; Wilson and Masel 2011). Selection on translation errors is for example strong enough to shape non-coding sequences beyond stop codons in *Saccharomyces cerevisiae* (Kosinski and Masel 2020).

Fitness effects in Neme et al. (2017) might not be directly caused by peptide properties alone but instead by the effect of both nucleotide and peptide properties on expression (Knopp and Andersson 2018), with lower expression being less harmful. For example, auto-downregulation at the mRNA level can cause significant difference in expression among peptides, despite identical promoters. However, the properties we find to be predictive, such as disorder and amino acid size, are not *a priori* related to auto-downregulation of mRNA in wild-type *E. coli*, making the latter an unlikely explanation for our findings.

Our findings are consistent with the hypothesis that peptides with low structural disorder tend to be harmful. We find that this effect is mediated by amino acid frequencies, with no additional contribution from amino acid ordering, at least none that could be picked up by use of the program IUPred. Disorder-promoting amino acids may help a peptide remain soluble even if unfolded. Small amino acids also tend to be benign, perhaps because they are hydrophobic enough to promote some amount of folding but flexible enough to avoid too much hydrophobic residue exposure.

Our findings suggest that the easiest way to avoid harm is through disorder and small size, but do not rule out other strategies that rely on capacity for folding. Indeed, BCS4, a *de novo* evolved protein in *Saccharomyces cerevisiae*, has a hydrophobic core and is capable of folding (Bungard, et al. 2017). Vakirlis et al. (2020) found that de novo proteins can emerge as transmembrane proteins, which need to be lipid soluble, presumably requiring different harm-avoidance strategies than peptides that are located in the cytosol.

The correlation between the extent to which an amino acid is enriched in young animal protein domains and its marginal fitness effect in random peptides in *E. coli* is intriguing, and consistent with a body of literature that *de novo* gene birth favors protein disorder. What is more, our ability to externally validate animal phylostratigraphy slopes against random peptides in *E. coli* provides additional support that these slopes represent more than mere bias, in contrast to suggests that all patterns are due to homology detection bias (Alba and Castresana 2007; Moyers and Zhang 2015, 2016). That is, if phylostratigraphy trends were due to an artifact such as homology detection bias, such an artifact would be unlikely to bias our random peptide analysis in the same direction.

Plants have different trends in amino acid frequencies as a function of sequence age than animals do, with young genes seeming to prefer readily available amino acids, rather than amino acids that promote ISD (James, et al. 2021). This could be because: 1) plants are less susceptible to harm from random peptides, 2) other properties, such as amino acid availability, drive the emergence of *de novo* genes in plants, or 3) the plant data lack the resolution needed to identify a correlation with the properties studied here. We do not have the ability to differentiate between these three possibilities here.

Nevertheless, our finding of consistency between what is benign in *E. coli* and what is benign in animals suggests the possibility of a deep concordance in what makes a peptide harmful between two apparently disparate branches of life. The forces that drive protein birth therefore appear to share a key similarity between bacteria and Animalia. Monod once suggested that what is true in *E. coli* must also be true in elephants; our work suggests that this may apply to the properties that tend to make peptides less harmful. To modify Monod’s famous quote, what is harmful in *E. coli* is also harmful in elephants, but not necessarily in eucalyptus.

A major idea in our understanding of proteins is that form – that is, the fold that is determined by the exact sequence of amino acids – determines function and thus fitness. However, for these random peptides in *E. coli*, the amino acid content but not the sequence in which they occur is the main determinant of benign vs harmful effects. Random peptides likely exist as a diverse ensemble of structural states, but the same is increasingly acknowledged to be true of functional proteins. While the ordering of amino acids in functional proteins no doubt plays a role, perhaps mere amino acid frequencies are also more important than once thought in this context too, especially in structurally disordered protein regions.

## Methods

### Data retrieval

Neme et al. (2017) performed seven experiments where *E. coli* lineages, each with a plasmid containing a unique random peptide, were grown and tracked using deep DNA sequencing. We downloaded sequencing counts from Dryad at http://dx.doi.org/10.5061/dryad.6f356, and obtained amino acid and nucleotide sequences directly from Rafik Neme. Experiment 7 was by far the largest with over 4 million reads, more than five times larger than the 2^nd^ largest experiment and over 1.2 million reads more than all other experiments combined. Experiment 7 contained all the peptides that the other six experiments classified as “increasing” or “decreasing,” and more. Small datasets from these other six experiments yield limited information because of the need to model changing mean fitness in a population, including not just the tracked lineages but also cells with an empty vector (see Estimating lineage fitness from random peptide sequencing counts section). We therefore chose to restrict our analysis to experiment 7.

Experiment 7 consists of the numbers of reads of each random peptide sequence in 5 replicate populations of *E. coli* at 4 time points. In principle, these independent measurements can simply be summed, allowing more precise fitness estimation than could be achieved from a single replicate. To first assess reproducibility, we estimated fitness for each replicate in isolation. Following Neme et al. (2017), we calculated fitness only when had ≥5 reads available, excluding cases in which all reads were on the same day. We grouped sequences into “clusters” to avoid pseudoreplication (see the “Clustering non-independent sequences” section below) and calculated the weighted mean fitness for each cluster. The five-choose-two pairs of replicates each shared between 612 and 624 clusters, with Pearson’s correlation coefficients ranging from 0.76 to 0.90 for fitness estimates of the same peptide clusters assessed in different replicates. We therefore summed across all five replicates to obtain a total number of reads for each polypeptide at each time point, and used these sums.

After summing across the first four replicates of experiment 7, we once again calculated fitness for peptides with ≥5 reads in our summed dataset, ending up with 1055 peptides out of over one million, grouped into 646 clusters. The amino acid sequences analyzed, the count data for each day and replicate of experiment 7, and the quantities we compute for the peptide, are available in our Supplemental Dataset S1, and the raw reads are available at the European Nucleotide Archive (ENA) under the project number PRJEB19640.

Neme et al. (2017) used this 5 read cutoff because it is not possible to infer fitness with any reasonable resolution for individual peptides with fewer than five reads. The dramatic nature of the data reduction from over a million peptides to only 646 clusters is unsurprising, firstly because each initial unique peptide was present in only one copy, and secondly because most peptides are likely deleterious. We note therefore that our analyzed subset of peptides with at least five reads are certainly non-lethal, and likely less deleterious than the average random peptide. Nonetheless, we achieved enough resolution to distinguish between more and less harmful peptides, with remarkably large effect sizes considering the restricted fitness range.

### Estimating lineage fitness from random peptide sequencing counts

The expected number of reads *λ*_*it*_ of peptide *i* at times *t*=1,2,3,4 was modeled as:

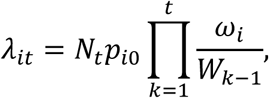

where *N*_*t*_ is the observed total number of reads, *p*_*i*0_ is the initial frequency of peptide *i* at the beginning of the experiment (prior to the round of selection used to produce the first measured timepoint *t* = 1), 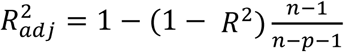 is the fitness of bacteria with peptide *i* at time *t* (i.e. their propensity to contribute to the next time point), and *W*_*k*_ is population mean fitness at time *k*, including bacteria containing empty vectors for which we have no direct count data.

The likelihoods of observed peptide counts were estimated from this expectation and two different error models. A Poisson distribution, which captures sampling error for count data given independence of each read, was used to generate our initial estimates of *p*_*io*_, *ω*_*i*_, and *W*_*k*_ (collectively yielding *λ*_*it*_) because it is analytically tractable. Under a Poisson error function, the likelihood of observing *n*_*it*_ reads of peptide *i* at time *t* is

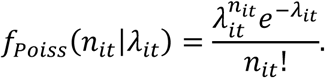

To relax the independence assumption to also capture variance inflation *κ* due to PCR amplification, we used a negative binomial distribution in the Polya form:

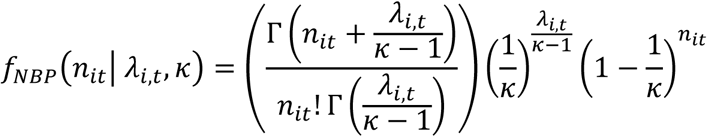

where Γ(⋅) is the gamma function. We used the initial estimates of *p*_*io*_, *ω*_*i*_, and *W*_*k*_ to numerically fit the negative binomial model. For the specifics of fitting the Poisson and negative binomial models, see Supporting Information. Weights were calculated, for use in downstream linear models, from this likelihood inference procedure, as the inverse of Fisher information (see Supporting Information). We use Fisher information to derive an estimate of the standard error from the curvature of the likelihood surface.

An existing software package for estimating lineage fitness from sequencing counts is Fit-Seq (Li, et al. 2018), which captures the amplification of PCR error through a more sophisticated distribution for the number of reads that is derived in the supplementary information of Levy et al. (2015). However, Fit-Seq assumes that mean fitness is a simple average of all measured lineages’ fitness, requiring all individuals to be tagged and measured. But Neme et al.’s (2017) experiment included lineages carrying an empty plasmid, i.e. with the selectable marker but no random peptide. Worse, the proportion of cells with an empty vector can be presumed to increase over time. In the absence of a reliable way to directly quantify cells with empty vectors, we instead consider mean population fitness over time to be a set of independent parameters to be fitted.

### Clustering non-independent sequences

Upon visual inspection, we found that some peptide sequences were extremely similar, with only one or two amino acid differences; these data points will not contain independent information about the relationship between sequence and fitness. To account for non-independence, we clustered peptides by their Hamming distance, and either took only the peptide whose fitness had the highest weight within its cluster, or took weighted means within clusters, or included cluster in our regression models as a random effect term. Single-link clustering with Hamming distance cutoffs of 6 to 29 amino acids all produced an identical set of 646 clusters for our 1051 peptides. The largest cluster had 228 random peptides, and the second largest had only 13. The vast majorities of clusters contained only 1 sequence (Dataset S1). A few peptides had mutations in their non-random regions; these mutations were counted in our Hamming distance measurements.

Such similar sequences are highly unlikely to arise by chance if the peptides were truly random; 20^50^ ≈ 10^65^ peptides are possible, far more than the ∼2 × 10^6^ observed. Because we analyze only peptides with at least 5 reads, replicated sequencing error is an unlikely cause. We see the same nearly-identical sequences appearing in every experimental replicate, suggesting either that mutations occurred during Neme et al.’s (2017) initial growth phase, or that the “random” peptides synthesized for the experiment are not entirely random. We note that construction of the “random” peptide library involved ligations of a smaller set of starting “seed” sequences, introducing non-randomness at this stage.

### Predictors of fitness

Neme et al. (2017) discarded all peptides with premature stop codons while producing the dataset we analyzed, so they are all exactly 65 amino acids long with 50 amino acids of random sequence. We therefore do not need to control for length, but do need to account for the mean fitness of a population that includes lineages expressing shorter peptides. An updated analysis was recently released that explored peptides of shorter, varying length (Castro and Tautz 2021).

#### GC content

Many amino acid sequences mapped to several possible nucleotide sequences, as part of the same problem of mutation or non-random construction discussed above. To calculate one GC content for each random peptide, we calculated a simple average of GC content across all the nucleotide sequences in the dataset that map to the peptide with the largest weight in the cluster, using only the random portion of the sequence.

#### Disorder

Protein disorder was measured using IUPred2 (Dosztányi, et al. 2005; Meszaros, et al. 2018) for amino acid sequences, and using disorder propensity (Theillet, et al. 2013) for individual amino acids. IUPred2 returns an ISD score between zero and one for each amino acid in a sequence, with higher scores indicating greater intrinsic disorder. To calculate an ISD score for each random peptide, we took the average of the scores for the whole sequence (i.e. including non-random parts). We used a square root transform because it produced a more linear relationship with fitness than no transform. All measurements referring to ISD or IUPred used IUPred2 except ΔISD, which used the original IUPred program – differences between the two are minimal (Meszaros, et al. 2018).

Disorder propensity gives each amino acid a score based on the frequency it is found in disordered proteins relative to ordered proteins (Theillet, et al. 2013). The disorder propensity score for a peptide was determined by averaging the disorder propensity scores for the amino acids in the random region. When we use the disorder propensity metric, we explicitly refer to it as “disorder propensity” and not as “ISD.”

#### Aggregation propensity

Tango (Fernandez-Escamilla, et al. 2004; Linding, et al. 2004; Rousseau, et al. 2006) returns an aggregation score for each amino acid in a sequence. At least five sequential amino acids with a score greater than or equal to five indicates an aggregation-prone region. We scored peptide aggregation propensity as the number of amino acids within regions scored as aggregation-prone, including contributions from non-random regions.

#### Solubility

CamSol (Sormanni, et al. 2015) returns a solubility score for each amino acid in a sequence, as well as a simple average of all scores for a sequence, which CamSol calls a “solubility profile.” We used the solubility profile of the full sequences, including non-random regions.

#### Amino acid frequencies

We counted frequencies among the 50 amino acids in the random portion of each peptide. The values for all the above predictors for each peptide are listed in Dataset S1.

### *E. coli* genome amino acid frequencies

Total amino acid frequencies in the *E. coli* genome were calculated from the K-12 reference proteome on UniProt (Bateman, et al. 2015), found at https://www.uniprot.org/proteomes/UP000000625. Excess/deficiency in frequency from that expected by codon number was determined by calculating the expected frequencies from codon number (e.g. 3/61 for isoleucine) and subtracting from the raw frequencies.

### Statistics

All statistical tests were carried out in R version 3.6.3 (R Core Team 2019), with figures generated using “ggplot2” (Wickham 2016). Weighted linear mixed models were implemented using the “lmer” function from the “lme4” package (Bates, et al. 2015), with cluster as a random effect. See Supporting Information for details, including justification of a log-transform for fitness. When R^2^ values were needed, we instead averaged peptides within the same cluster into a combined datapoint, allowing us to avoid the use of a random effect term. We calculated adjusted R^2^ values using the base R “lm” function. Adjusted R^2^ is a modification of R^2^ to penalize additional predictors, and is calculated using the formula:

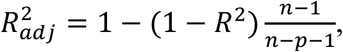

where *n* are the number of data points and *p* are the number of predictors. Raw P-values are reported unless otherwise noted, i.e. without correction for multiple comparisons.

## Data and code availability

All code and supplemental data files are available on GitHub at https://github.com/MaselLab/RandomPeptides. The data originally released in conjunction with the publication of Neme et al. (2017) can be found at Dryad http://dx.doi.org/10.5061/dryad.6f356, and the raw reads are available at the European Nucleotide Archive (ENA) under the project number PRJEB19640.

## Acknowledgements

This work was supported by the John Templeton Foundation (39667, 60814) and the National Institutes of Health (GM-104040, T32GM-008659, T32GM-084905). We thank Rafik Neme and Diethard Tautz for sharing their data with us and for graciously answering all our questions regarding their analyses, Dvir Schirman and Tzachi Pilpel for sharing their data with us, Joe Watkins for helpful discussions about our likelihood estimation procedure, and Catherine Weibel for driving the GC content analysis forward.

## Notes

https://github.com/MaselLab/RandomPeptides

### Competing Interest Statement

The authors have declared no competing interest.

### Summary of Updates

Added a new figure. Minor changes to clarify methods. A bug was caught that had very minor effects on the numbers throughout.

