## Supplemental information for "Random peptides rich in small and disorder-promoting amino acids are less likely to be harmful"

### Supplementary Text

#### Estimating lineage fitness from random peptide sequencing counts

**Fitting the Poisson model.** We obtain analytical expressions for the maximum likelihood values of the peptide-specific parameters, namely $p_{i0}$ and $\omega_i$, while holding mean population fitness, $W_k$, fixed, as well as for the maximum likelihood value of $W_k$ while holding $p_{io}$and $\omega_i$ fixed. As discussed above, $W_k$ is not a weighted mean of $\omega_i$, because $\omega_i$ estimates are not produced for all lineages – many go rapidly extinct, and some also have an empty vector that does not express a random peptide, and therefore do not appear in read counts. We alternate between holding $W_k$ fixed and holding $p_{io}$and $\omega_i$ fixed until convergence of our estimates.

To estimate $p_{i0}$ and $\omega_i$ while holding $W_k$ fixed, we need to maximize one likelihood function for each peptide:

$$L(\lambda_{it}|\boldsymbol{n_{it}}) = \prod_t \frac{\lambda_{it}^{n_{it}} e^{-\lambda_{it}}}{n_{it}!}.$$

We take the logarithm

$$\ln\big(L(\lambda_{it}|\boldsymbol{n_{it}})\big) = \sum_t (n_{it} \ln \lambda_{it}) - \sum_t \lambda_{it} - \sum_t \ln(n_{it}!),$$

and then solve for a maximum by considering the set of critical points. First, we take the derivative with respect to $p_{i0}$:

$$\frac{\partial}{\partial p_{i0}} \ln\big(L(\lambda_{it}|\boldsymbol{n_{it}})\big) = \sum_t \frac{n_{it}}{p_{i0}} - \sum_t \left[N_t \prod_{k=1}^{t} \frac{\omega_i}{W_{k-1}}\right].$$

Setting this equal to zero, we simplify to obtain

$$p_{i0} \sum_t \left[N_t \prod_{k=1}^{t} \frac{\omega_i}{W_{k-1}}\right] = \sum_t n_{it},$$

yielding

$$p_{i0} = \frac{\sum_t n_{it}}{\sum_t \left[N_t \prod_{k=1}^{t} \frac{\omega_i}{W_{k-1}}\right]}.$$

Similarly, by taking the derivative of $\ln\big(L(\lambda_{it}|\boldsymbol{n_{it}})\big)$ with respect to $\omega_i$ and solving again for $p_{i0}$, we obtain

$$p_{i0} = \frac{\sum_t n_{it} t}{\sum_t \left[N_t t \prod_{k=1}^{t} \frac{\omega_i}{W_{k-1}}\right]}.$$

Thus, with our two equations for $p_{i0}$, we have

$$p_{i0} = \frac{\sum_t n_{it}\, t}{\sum_t \left[N_t t \prod_{k=1}^{t} \frac{\omega_i}{W_{k-1}}\right]} = \frac{\sum_t n_{it}}{\sum_t \left[N_t \prod_{k=1}^{t} \frac{\omega_i}{W_{k-1}}\right]}.$$

Cross multiplying these two terms gives us

$$\left[\sum_t N_t \prod_{k=1}^{t} \frac{\omega_i}{W_{k-1}}\right] \sum_t n_{it} t = \left[\sum_t N_t t \prod_{k=1}^{t} \frac{\omega_i}{W_{k-1}}\right] \sum_t n_{it},$$

and thus

$$0 = \left[\sum_t N_t \prod_{k=1}^{t} \frac{\omega_i}{W_{k-1}}\right] \sum_t n_{it} t - \left[\sum_t N_t t \prod_{k=1}^{t} \frac{\omega_i}{W_{k-1}}\right] \sum_t n_{it}.$$

We pull out the sum and simplify as such:

$$0 = \sum_t \left[\left[N_t \prod_{k=1}^{t} \frac{\omega_i}{W_{k-1}}\right] \left(\sum_k n_{ik} k - t \sum_k n_{ik}\right)\right].$$

Note that we have changed index on the inner sums to avoid summing over the same index on the inner and outer sums. After reorganizing for clarity,

$$0 = \sum_t \left[\left(\sum_k n_{ik} k - n_{ik} t\right) \left[N_t \prod_{k=1}^{t} \frac{\omega_i}{W_{k-1}}\right]\right]$$

$$= \sum_t \left[\left(\sum_k (k-t) n_{ik}\right) \left[N_t \prod_{k=1}^{t} \frac{\omega_i}{W_{k-1}}\right]\right].$$

As the expression is set equal to zero, we can consider the additive inverse of the right-hand side as follows (multiplying through by -1):

$$0 = \sum_t \left[\left(\sum_k (t-k) n_{ik}\right) \left[N_t \prod_{k=1}^{t} \frac{\omega_i}{W_{k-1}}\right]\right].$$

Now if we let

$$\mathrm{C_t} = \left(\sum_k (t-k) n_{ik}\right) \left[N_t \prod_{k=1}^{t} \frac{1}{W_{k-1}}\right],$$

we find our expression is a polynomial of the form

$$0 = \sum_t C_t \omega_i^t.$$

Noting that all terms have time $t \geq 1$, we find $\omega_i = 0$ to be a trivial solution. However, $\omega_i = 0$ cannot be a solution as the Poisson model requires a positive rate parameter. We then solve for $\omega_i$ numerically and find the corresponding $p_{i0}$ using either of the equations we derived earlier, and then select our estimates as the pair that maximizes the likelihood function.

For the next part of our iterative procedure, we use the solutions above as parameters for our model and estimate the values of the mean fitness trajectory $\{W_t\}_{0 \leq t \leq 3}$. Here we must calculate the likelihood across the entire population at each time point instead of a single peptide at a time:

$$L(\lambda_{it} | n_{it}) = \prod_i \prod_t \frac{\lambda_{it}^{n_{it}} e^{-\lambda_{it}}}{n_{it}!}.$$

Taking the logarithm, we obtain

$$\ln\big(L(\lambda_{it} | \boldsymbol{n_{it}})\big) = \sum_i \left(\sum_t n_{it} \ln \lambda_{it} - \lambda_{it} - \ln(n_{it}!)\right).$$

The process of getting these estimators will be recursive, as the mean fitness correction at time $t$ depends on time $t-1$, and $t-1$ on time $t-2$ etc. We therefore start at the final time $\tau = 3$, which is the only time $W_\tau$ appears. Taking the derivative with respect to the final time $W_\tau$ of the log-likelihood function gives

$$\frac{\partial}{\partial W_\tau}\ln\big(L(\lambda_{it}|\boldsymbol{n_{it}})\big) = \frac{1}{W_\tau^2}\sum_i\left[N_\tau p_{i0}\prod_{k=1}^{\tau+1}\frac{\omega_i}{W_{k-1}}\right] - \sum_i\left(\frac{n_{i\tau}}{W_\tau}\right)$$

Setting this equal to zero to solve for the critical value gives

$$\sum_i\left(\frac{n_{i\tau}}{W_\tau}\right) = \frac{1}{W_\tau^2}\sum_i\left[N_\tau p_{i0}\,\omega_i\prod_{k=1}^{\tau}\frac{\omega_i}{W_{k-1}}\right],$$

and with some simplification we find

$$W_\tau\sum_i n_{i\tau} = \sum_i\left[N_\tau p_{i0}\,\omega_i\prod_{k=1}^{\tau}\frac{\omega_i}{W_{k-1}}\right],$$

which provides

$$W_\tau = \frac{N_\tau}{\sum_i n_{i\tau}}\frac{\sum_i p_{i0}\,\omega_i^\tau}{\prod_{k=1}^{\tau} W_{k-1}}.$$

Prior to generalizing, we next demonstrate the specific derivation of $W_{\tau-1}$. Taking the logarithm, we again have

$$\ln\big(L(\lambda_{it}|\boldsymbol{n_{it}})\big) = \sum_i\left(\sum_t n_{it}\ln\lambda_{it} - \lambda_{it} - \ln n_{it}!\right)$$

After taking the derivative with respect to $W_{\tau-1}$ we obtain additional terms dependent on $W_{\tau-1}$.

$\frac{\partial}{\partial W_{\tau-1}}\ln\big(L(\lambda_{it}|\boldsymbol{n_{it}})\big) = \sum_i\left[-\left(\frac{n_{i(\tau-1)}}{W_{\tau-1}}\right) + \left[N_{\tau-1}p_{i0}\prod_{k=1}^{\tau}\frac{\omega_i}{W_{k-1}}\right]\left(\frac{1}{W_{\tau-1}}\right)\right] + \sum_i\left[-\left(\frac{n_{i\tau}}{W_{\tau-1}}\right) + \left[N_\tau p_{i0}\prod_{k=1}^{\tau+1}\frac{\omega_i}{W_{k-1}}\right]\left(\frac{1}{W_\tau W_{\tau-1}}\right)\right]$

Now just consider the second summation

$$= \sum_i\left[-\left(\frac{n_{i\tau}}{W_{\tau-1}}\right) + \left[N_\tau p_{i0}\prod_{k=1}^{\tau+1}\frac{\omega_i}{W_{k-1}}\right]\left(\frac{1}{W_\tau W_{\tau-1}}\right)\right]$$

$$= \left[N_\tau\left(\frac{1}{W_\tau W_{\tau-1}}\right)\prod_{k=1}^{\tau+1}\frac{\omega_i}{W_{k-1}}\right]\left[\sum_i p_{i0}\,\omega_i^\tau - \frac{1}{W_{\tau-1}}\sum_i n_{i\tau}\right]$$

Substituting our earlier result that

$$W_\tau = \frac{N_\tau}{\sum_i n_{i\tau}}\frac{\sum_i p_{i0}\,\omega_i^\tau}{\prod_{k=1}^{\tau} W_{k-1}},$$

and through cancelation we obtain

$$\frac{1}{W_{\tau-1}}\sum_i n_{i\tau} - \frac{1}{W_{\tau-1}}\sum_i n_{i\tau} = 0.$$

Thus, we come to an expression with the same structure as before with $W_\tau$

$$\frac{\partial}{\partial W_{\tau-1}}\ln\big(L(\lambda_{it}|\boldsymbol{n_{it}})\big) = \frac{1}{W_{\tau-1}^2}\sum_i\left[N_{\tau-1}p_{i0}\prod_{k=1}^{\tau}\frac{\omega_i}{W_{k-1}}\right] - \sum_i\frac{n_{i(\tau-1)}}{W_{\tau-1}}.$$

Setting this equal to zero to solve for the critical value and rearranging terms gives

$$\sum_i \frac{n_{i(\tau-1)}}{W_{\tau-1}} = \frac{1}{W_{\tau-1}^2}\sum_i \left[N_{\tau-1}p_{i0}\prod_{k=1}^{\tau}\frac{\omega_i}{W_{k-1}}\right],$$

and with some simplification

$$W_{\tau-1}\sum_i n_{i(\tau-1)} = \sum_i \left[N_{\tau-1}p_{i0}\omega_i\prod_{k=1}^{\tau}\frac{\omega_i}{W_{k-1}}\right],$$

which provides

$$W_{\tau-1} = \frac{N_{\tau-1}}{\sum_i n_{i(\tau-1)}}\frac{\sum_i p_{i0}\omega_i^{\tau-1}}{\prod_{k=1}^{\tau} W_{k-1}}.$$

Applying the same method to $W_{\tau-2}$, and so on, using the same strategy of substitution, cancels related terms for $t > \tau - 2$ and gives a solution of the same form as the one above.

To generalize this, let $\rho_t = \frac{\sum_i n_{it}}{N_t}$ be the observed fraction of total reads represented by the peptides studied in detail, and let $\phi_t = \sum_i p_{i0}\omega_i^t$. We find that all estimators of mean fitness correction have the form

$$W_t = \frac{\rho_{t-1}}{\rho_t}\frac{\phi_t}{\phi_{t-1}} = \left[\frac{\phi_t}{\phi_{t-1}}\right] / \left[\frac{\rho_t}{\rho_{t-1}}\right]$$

and thus, as would be found through repeating the process of substitution above,

$$W_1 = \frac{\phi_1}{\rho_1}.$$

These are the sums of the expected changes in frequency divided by the sum of the actual total proportion observed at time t.

**Fitting the Negative Binomial model.** The following is the likelihood function of our Negative Binomial model:

$$L(\lambda_{it}|\boldsymbol{n_{it}}) = \prod_t \left[\left(\frac{\Gamma\left(n_{it}+\frac{\lambda_{it}}{\kappa-1}\right)}{\Gamma\left(\frac{\lambda_{it}}{\kappa-1}\right)n_{it}!}\right)\left(\frac{1}{\kappa}\right)^{\frac{\lambda_{it}}{\kappa-1}}\left(1-\frac{1}{\kappa}\right)^{n_{it}}\right].$$

As stated, unlike with the Poisson model, we fit our negative binomial model numerically. The general form with respect to $p_{i0}$ and $\omega_i$ would require us to find the solutions to the following expressions

$$0 = \sum_t \Psi\left(n_{it} + \frac{\lambda_{it}}{\kappa-1}\right) - \Psi\left(\frac{\lambda_{it}}{\kappa-1}\right) - \frac{\partial}{\partial\theta}\lambda_{it}\left(\frac{\ln(\kappa)}{\kappa-1}\right)$$

$$\theta: p_{i0}, \omega_i, \{W_t\}_{0\le t\le 3}$$

and for the $\kappa$ inflation parameter we have

$$0 = \sum_t \Psi\left(n_{it} + \frac{\lambda_{it}}{\kappa-1}\right) - \Psi\left(\frac{\lambda_{it}}{\kappa-1}\right) - \lambda_{it}\left(\frac{\kappa - \kappa\ln(\kappa) - 1}{\kappa(\kappa-1)^2}\right) + \frac{n_{it}}{\kappa(\kappa-1)}$$

where $\Psi(\cdot)$ is the di-gamma function, the first derivative of the log-gamma function. As the di-gamma function has no closed form, we did not use the equation above, but proceeded numerically by implementing the FindMaximum function in Mathematica version 11.1.1 (Wolfram Research 2017), optimizing our parameters stepwise over a select set of them at a time, fixing all others as constant.

Our procedure begins with obtaining maximum likelihood estimates for $W_t$, and for $\omega_i$ and $p_{i0}$ across all peptides under the Poisson model to serve as initial values in our maximization procedure. Our Poisson initial estimates are equivalent to a negative binomial model in which we take the limit as the variance inflation $\kappa$ approaches 1. Using $\omega_i$ and $p_{i0}$ for each peptide population in our Negative Binomial model, we next estimate the variance inflation $\kappa$ for the first time. We then repeat our estimation of $\omega_i$ and $p_{i0}$, followed by $W_t$, and then again $\kappa$, continuing until the estimates converge.

In our Mathematica implementation of the process described above, we used the default settings for the FindMaximum function, using each updated estimate as the initial values for the next iteration of our maximization while searching within predefined bounded intervals. We estimated $p_{i0}$ in the interval (0,1) given the assumption that all peptides were present at the beginning of the experiment. Our measure of peptide fitness contribution $\omega_i$ was estimated from the interval (0,20], given that we should have non-zero expectation as required with the Poisson model, as well as to reduce computation time by limiting our search interval with an upper bound greater than all previously seen Poisson estimates. Similar reasoning applies to the restriction of the search for the mean fitness trajectory within the range of (0, 5).

All our fitness effect estimates for the random peptides using the negative binomial model are listed in supplemental Dataset S1.

#### Calculating weights for our fitness estimates.

Peptides vary greatly with their total sequencing counts; some have thousands of counts, while some have as few as five. Count variation introduces heteroscedasticity into our fitness estimates, which we account for by weighting our estimates. To obtain the weights used in our regression models, we approximate the variance of our estimators with the observed Fisher information. To estimate the Fisher information in $\omega_i$, given data $\{n_i\}$ drawn from a sampling process

$$I(\omega_i) = -\mathbb{E}_\omega\left[\frac{\partial^2}{\partial\omega_i^2} ln(L_x(\{n_i\}|\omega_i))\right],$$

we calculated observed Fisher information (Efron and Hinkley 1978), i.e. the Fisher information in our estimate $\widehat{\omega}_i$, defined as

$$I(\widehat{\omega}_i) = -\frac{\partial^2}{\partial\omega_i^2} ln\big(L_x(\{n_i\}|\widehat{\omega}_i)\big).$$

We approximate reads at different timepoints as being independently sampled, i.e. we neglect the intrinsic stochastic amplification by which a peptide that is oversampled at timepoint 2 will have a frequency advantage in timepoint 3. This yields

$$I(\widehat{\omega}_i) = -\sum_{t=1}^{k}\frac{\partial^2}{\partial\omega_i^2} ln\big(f_{NBP}(n_{it}|\widehat{\omega}_i)\big)$$

where $\widehat{\omega}_i$, $\kappa$, and $p_{i0}$ determine the expectation $\lambda_{i,t}$ in our previous expression for $f_{NBP}$. Thus, the observed Fisher information for the Negative Binomial model is

$$I(\widehat{\omega}_i) = -\sum_{t=1}^{k}\left[\Psi'\left(n_{it}+\frac{\lambda_{it}}{\kappa-1}\right)-\Psi'\left(\frac{\lambda_{it}}{\kappa-1}\right)-\frac{\partial^2}{\partial\omega^2}\lambda_{it}\left(\frac{\ln(\kappa)}{\kappa-1}\right)\right]$$

where $\Psi'(\cdot)$ indicates the first derivative of the digamma function (the second derivate of the log-gamma function).

The weights on the fitness estimates for each peptide are listed in supplemental Dataset S1.

#### Improved power for detecting candidate beneficial peptides

The statistical analysis performed by Neme et al. (2017) focused on the 1082 peptides with at least five sequencing reads. Out of these 1082 peptides, Neme et al. (2017) found that 277 were increasing, while 436 were decreasing. Using fitness values inferred as described above, we find statistical support for increases in frequency (i.e. fitness – 1.96 × standard error > 1) for 475 peptides (Dataset S1). We exclude 22 of the previously reported 277, at least some of which arose from a bug we located in Neme et al.'s

(2017) pipeline. 509 peptides had statistical support for decreases in frequency, with support lost for 9 of the previously reported 436 deleterious peptides.

#### Regression models

We used weighted linear mixed models with sequence cluster as a random effect, implemented in R using the "lmer" function from the "lme4" package (Bates et al. 2015), to determine which factors predict the estimated fitness of each peptide. Weights were based on the inverse of Fisher information during the maximum likelihood inference of each estimated fitness value (see "Calculating weights for our fitness estimates" section above). After finding that amino acid composition was by far the strongest predictor of fitness (see Results), we tested other predictors to see if they significantly improved the fit over an amino acid composition-only model. We do not include an intercept in the model; because one amino acid is redundant, a model with 19 amino acids and an intercept is equivalent to a model with 20 amino acids and no intercept. We tested IUPred-predicted disorder, Tango-predicted aggregation propensity, and CamSol-predicted solubility. None of the additional predictors improved the model beyond the base model of amino acid composition alone (supplemental Table S1).

We did not transform fitness values, but we did weight by $1/_{var}$, where $var$ was estimated using observed Fisher information (see "Calculating weights for our fitness estimates section above). We observed some mild systematic heteroscedasticity (supplemental fig. S2A), but transforming for normality to remove the skew in fitness values would makes matters worse rather than better (supplemental fig. S2B).

#### Calculating each amino acid's marginal effect on fitness

The marginal effect of x on y is the change in y when only x varies. But each of our random sequences is exactly 50 amino acids long, so adding one amino acid of any type must also result in losing an amino acid of a different type, whose identity must be chosen from an appropriate probability distribution. For each of the 646 clusters, we take the weighted mean of each amino acid count in the random portion of the peptide, and then we calculate the amino acid counts $AA_i$ as a simple sum across these 646 measures. The marginal fitness effect can then be calculated from the set of fixed effects $\beta$ in our weighted linear mixed model with cluster as a random effect and only amino acid frequencies as fixed effects. The marginal fitness effect of amino acid $i$ is then

$$\beta_i - \sum_{j \neq i} \left( \frac{AA_j}{\sum_{k \neq i} AA_k} \right) \beta_j$$

Here, $\frac{AA_j}{\sum_{k \neq i} AA_k}$ is the frequency of amino acid $j$ in the data set after amino acid $i$ is excluded, and represents the probability of losing an amino acid of type $j$. We use the effect values $\beta$ from our weighted linear mixed model (i.e. not collapsing clusters into summaries) where just the 20 amino acid counts were used to predict $\log_2$ fitness.

The variance of the marginal effect of each amino acid $i$ is $\mathrm{var}\left(\beta_i - \sum_{j \neq i} \left( \frac{AA_j}{\sum_{k \neq i} AA_k} \right) \beta_j \right)$. Expanding this expression generates many covariance terms because the $\beta'$s are not independent. The expansion can be obtained by rewriting

$$\beta_i - \sum_{j \neq i} \left( \frac{AA_j}{\sum_{k \neq i} AA_k} \right) \beta_j = \sum_l C_l \beta_l \,,$$

where the coefficients

$$C_l = \begin{cases} 1 & l = i \\ -\frac{AA_l}{\sum_{k \neq i} AA_k} & l \neq i, \end{cases}$$

are those of each $\beta$ in the original expression $\beta_i - \sum_{j \neq i} \left( \frac{AA_j}{\sum_{k \neq i} AA_k} \right) \beta_j$. Thus,

$$\mathrm{var}\left( \beta_i - \sum_{j \neq i} \left( \frac{AA_j}{\sum_{k \neq i} AA_k} \right) \beta_j \right) = \mathrm{var}\left( \sum\nolimits_l C_l \beta_l \right)$$

and we apply the property,

$$\mathrm{var}\left( \sum\nolimits_l C_l \beta_l \right) = \sum\nolimits_l C_l^2 \mathrm{var}(\beta_l) + 2 \sum\nolimits_{l<m} C_l C_m \mathrm{cov}(\beta_l, \beta_m).$$

We use the right-hand side to calculate the variance of the marginal effect for each amino acid by substituting in for the $C$'s and the variances and covariances of the $\beta$'s. Note that error in $AA_i$ is negligible compared to error in $\beta_i$, so we do not consider its variance. Variances and covariances were obtained by calling R's "vcov" function on the lmer mixed model object for our weighted mixed model where just the 20 amino acid frequencies were used to predict fitness. Marginal effects with standard errors for each amino acid are shown in supplemental Table S2.

#### Supplemental Figures, Tables, and Datasets

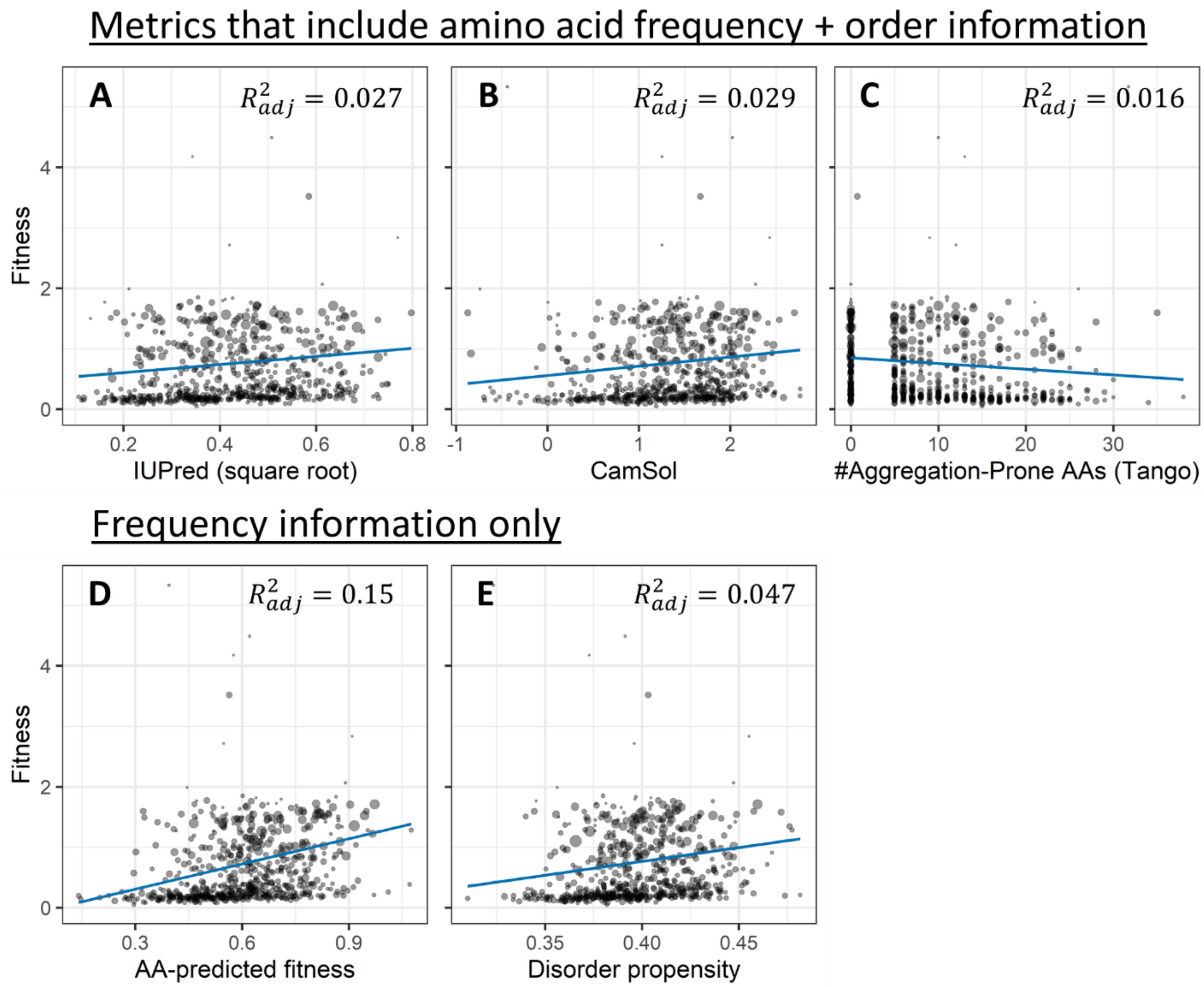

**Figure S1**
Zoomed out versions of fig. 1 plots, to include the seven clusters with fitness greater than 2. The figure details are the same as in fig. 1.

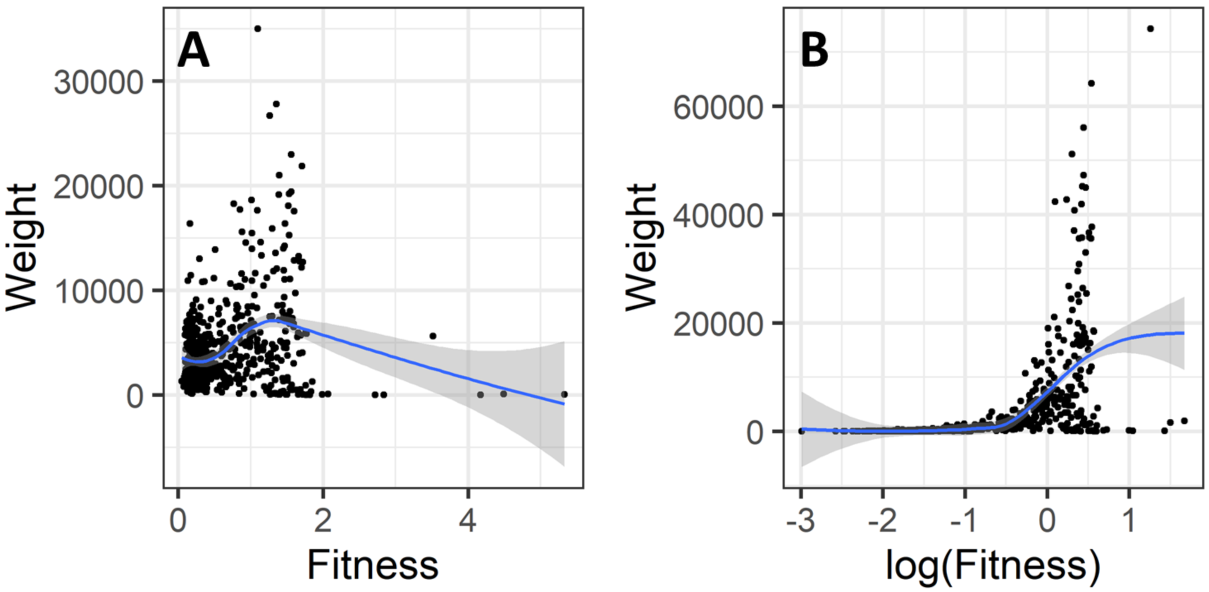

**Figure S2**

Untransformed fitness is appropriate for use in regression models. Each point represents a cluster (n = 646, see “Clustering non-independent sequences” in the Methods), showing only the highest-weighted peptide within each cluster. Weights are calculated as $^1/_{var}$, where observed Fisher information is used to estimate variance (see “Calculating weights for our fitness estimates” section above). In log space, the second derivative with respect to $\log(\omega)$ rather than $\omega$ yields a different weight. A skewed distribution with a small number of high fitness values suggests a log transform, which would yield a more normal distribution. But untransformed fitness and associated weights (A) show mild systematic heteroscedasticity with little correlation with fitness, while log fitness (B) shows a stronger dependence of weight on fitness. This suggests that a model using untransformed fitness is most appropriate. The blue line is a loess curve with confidence intervals in grey.

#### Table S1

| PREDICTOR OF $\mathbf{Ln}(\boldsymbol{Fitness})$ | $\boldsymbol{\beta}$ | STANDARD ERROR | P-VALUE |
|---|---|---|---|
| Ile | -0.05864 | 0.016283 | N/A |
| Phe | -0.02159 | 0.015093 | N/A |
| His | -0.01937 | 0.0178 | N/A |
| Met | -0.01798 | 0.022434 | N/A |
| Asn | -0.00938 | 0.022098 | N/A |
| Tyr | -0.00806 | 0.019885 | N/A |
| Lys | -0.00602 | 0.02065 | N/A |
| Val | -0.00474 | 0.009331 | N/A |
| Leu | -0.0038 | 0.008792 | N/A |
| Glu | -0.00277 | 0.016185 | N/A |
| Arg | 0.003363 | 0.007974 | N/A |
| Cys | 0.007966 | 0.013214 | N/A |
| Thr | 0.010522 | 0.01372 | N/A |
| Trp | 0.021886 | 0.017064 | N/A |
| Gly | 0.028399 | 0.008036 | N/A |
| Asp | 0.030869 | 0.017118 | N/A |
| Gln | 0.033101 | 0.017284 | N/A |
| Ser | 0.038431 | 0.008247 | N/A |
| Pro | 0.044911 | 0.012225 | N/A |
| Ala | 0.045448 | 0.009858 | N/A |
| $\mathrm{Tango}$ | -0.0029 | 0.0032 | 0.3 |
| $\sqrt{\mathrm{ISD}}$ | -0.96 | 0.74 | 0.2 |
| $\mathrm{CamSol}$ | 0.09 | 0.077 | 0.2 |

Mixed model predictors of lineage fitness. P-values come from a likelihood ratio test of either dropping the amino acid from a model with all amino acids predicting fitness or adding the non-amino acid predictor to the same model. β coefficients for amino acids come from a model with only the 20 amino acids predicting fitness, and β coefficients for non-amino acid predictors come from adding the predictor to the 20 amino acid-only model. Note that while the beta coefficient for ISD appears large compared to other coefficients, this is because ISD as assessed by an IUPred score is a number between 0 and 1, whereas amino acids are count data and hence integers – any comparison must involve the product of the coefficient and a predictor score of appropriate magnitude. IUPred, Tango, and CamSol scores are not predictive ($P>0.05$) once amino acid frequencies are controlled for. Note that our hypotheses about the β coefficients of individual amino acids pertain to their values relative to one another rather than relative to zero, so we report their p-values for inclusion in the model as “N/A.”

#### Table S2

| AMINOACID | EFFECT ON GENOTYPE FREQUENCY / CYCLE | STD.ERR |
|---|---|---|
| **ILE** | -0.07297 | 0.0168 |
| **PHE** | -0.03483 | 0.01561 |
| **HIS** | -0.03217 | 0.018204 |
| **MET** | -0.03057 | 0.022813 |
| **ASN** | -0.02183 | 0.022479 |
| **TYR** | -0.0206 | 0.020335 |
| **VAL** | -0.01846 | 0.010228 |
| **LYS** | -0.01846 | 0.021054 |
| **LEU** | -0.01751 | 0.009677 |
| **GLU** | -0.01531 | 0.016679 |
| **ARG** | -0.0098 | 0.008953 |
| **CYS** | -0.00431 | 0.01383 |
| **THR** | -0.00163 | 0.014308 |
| **TRP** | 0.010089 | 0.017557 |
| **GLY** | 0.018508 | 0.009103 |
| **ASP** | 0.019292 | 0.017576 |
| **GLN** | 0.021556 | 0.017724 |
| **SER** | 0.028946 | 0.009051 |
| **PRO** | 0.034615 | 0.012885 |
| **ALA** | 0.036672 | 0.010825 |

Marginal effect of replacing one amino acid (with fitness estimated in Table S1) with a randomly chosen amino acid. Available for download at https://github.com/MaselLab/RandomPeptides/blob/master/Data/supplemental_table_2.tsv

#### Dataset S1

Table of all predictors, sequences, and sequencing counts for all the peptides analyzed in our study. The first column, “PeptideID,” is the peptide unique ID and matches those used by Neme et al. (2017). “AASeq” is the column for the amino acid sequence of the random peptide; “Cluster” is for the cluster assigned to non-independent sequences; “Fitness.nb” and “Weight.nb” are for our fitness estimates and weights on those estimates, respectively, for random peptides from our negative binomial model. “PredHel” is a binary where a “1” indicates that the peptide has a predicted trans-membrane helix from TMHMM (Krogh et al. 2001), and a “0” indicates no transmembrane helix was predicted. The predictors we examined (e.g. IUPred2 (Dosztányi et al. 2005; Meszaros et al. 2018), CamSol (Sormanni et al. 2015), GC content) are also listed separately in their own columns with appropriate names. Columns for counts of each amino acid per peptide are labeled by their respective amino acid’s three letter name. Sequencing count data is labeled in a “dX-rY” format, where “dX” is the day, and “rY” is the replicate (e.g. d1-r1 is the sequencing counts for the first replicate in the first day). The table is too large to display here and may be downloaded at https://github.com/MaselLab/RandomPeptides/blob/master/Data/supplemental_dataset_1.tsv.

Wolfram Research, Inc. 2017. Mathematica. Version 11.1.1. Champaign, Illinois: Wolfram Research, Inc.
